# Genome analysis reveals evolutionary mechanisms of adaptation in systemic dimorphic fungi

**DOI:** 10.1101/199596

**Authors:** José F. Muñoz, Juan G. McEwen, Oliver K. Clay, Christina A. Cuomo

## Abstract

Dimorphic fungal pathogens cause a significant human disease burden and unlike most fungal pathogens affect immunocompetent hosts. To examine the origin of virulence of these fungal pathogens, we compared genomes of classic systemic, opportunistic, and non-pathogenic species, including *Emmonsia* and two basal branching, non-pathogenic species in the Ajellomycetaceae, *Helicocarpus griseus* and *Polytolypa hystricis*. We found that gene families related to plant degradation, secondary metabolites synthesis, and amino acid and lipid metabolism are retained in *H. griseus* and *P. hystricis*. While genes involved in the virulence of dimorphic pathogenic fungi are conserved in saprophytes, changes in the copy number of proteases, kinases and transcription factors in systemic dimorphic relative to non-dimorphic species may have aided the evolution of specialized gene regulatory programs to rapidly adapt to higher temperatures and new nutritional environments. Notably, both of the basal branching, non-pathogenic species appear homothallic, with both mating type locus idiomorphs fused at a single locus, whereas all related pathogenic species are heterothallic. These differences revealed that independent changes in nutrient acquisition capacity have occurred in the Onygenaceae and Ajellomycetaceae, and underlie how the dimorphic pathogens have adapted to the human host and decreased their capacity for growth in environmental niches.

## INTRODUCTION

Among millions of ubiquitous fungal species that pose no threat to humans, a small number of species cause devastating diseases in immunocompetent individuals. Notably, many human pathogenic species are ascomycetes in the order Onygenales, which comprises dermatophytes (e.g. *Trichophyton* spp.), the spherule-forming dimorphic pathogen *Coccidioides*, and yeast-forming dimorphic pathogens *Histoplasma*, *Paracoccidioides*, and *Blastomyces*. Dimorphism is a specialized morphogenetic adaptation allowing both growth in the environment and colonization of a host and is critical for the lifecycle of dimorphic fungal pathogens ^1^. In yeast-forming pathogens, dimorphism has been broadly defined as the ability to generate both yeast (e.g. blastoconidia) in the host or at 37 °C, and mycelia (e.g. conidia-producing or other hyphae) in the environment ^1^. During fungal evolution, dimorphism has arisen independently multiple times in saprophytic fungi, as is manifested in the wide distribution of dimorphic species across the Ascomycota, Basidiomycota and Zygomycota phyla ^2^. Notably, the dimorphic fungi in the order Onygenales are primary pathogens causing systemic mycosis in healthy humans, and collectively cause over 650,000 new infections a year in the United States alone ^3^. Dimorphic fungi can persist as latent infections in tens of millions of people worldwide and may reactivate when the host becomes immune-deficient; symptoms of active infection include pneumonia, acute respiratory distress syndrome, and disseminated disease, which can affect multiple organ systems ^1^.

The largest cluster of thermally dimorphic human pathogenic fungi belongs to the onygenalean family Ajellomycetaceae, which includes *Histoplasma*, *Paracoccidioides*, and *Blastomyces*. In addition to these medically important genera, the Ajellomycetaceae family also includes more rarely observed pathogenic species including *Emmonsia parva* and *Ea. crescens*, which undergo a thermal dimorphic transition to produce adiaspores rather than yeast and cause adiospiromycosis sporadically in humans ^4,5^. This group also includes *Lacazia loboi*, the yeast-like etiological agent of lobomycosis ^6^. Of public health concern, recent reports have documented the worldwide emergence of yeast-forming Ajellomycetaceae that cause systemic mycoses predominantly in immunocompromised patients, often with high case-fatality rates ^7,8^. Morphological and phylogenetic analyses were combined to demarcate species boundaries within the Ajellomycetaceae with clinical significance, which led to a proposed revision of the taxonomy; this included addition of the new genus *Emergomyces*, which includes the new species *Es. africanus* and *Es. orientalis*, the description of the new species *Blastomyces percursus*, and ongoing efforts to define additional *Emmonsia*-like and *Blastomyces*-like species ^8–10^.

Dimorphic species within the Ajellomycetaceae are typically restricted to specific ecological niches and geographical regions. For example, while *H. capsulatum* and *Es. pasteurianus* are considered to have worldwide distribution, *B. dermatitidis*, *P. brasiliensis*, and *Es. africanus* are restricted to North America, Latin America and South Africa, respectively. Non-dimorphic species have also been documented as early diverging Ajellomycetaceae, including *Emmonsiellopsis*, *Helicocarpus griseus* and *Polytolypa hystricis*, neither of which has been associated with disease in mammals. These species are found in environmental samples and do not appear to undergo a thermally regulated dimorphic transition ^11–13^.

We hypothesized that such phenotypic differences within the Ajellomycetaceae could be associated with gene family expansions or contractions, as a consequence of gene duplication and gene loss events, and that species comparison might reveal evolutionary mechanisms of adaptation in the systemic dimorphic fungi. Previous comparative and population genomic studies within Ajellomycetaceae had found evidence of gene family contractions and expansions associated with virulence ^5,14,15^. Other Onygenales genera outside the Ajellomycetaceae represented by published genome sequences include *Coccidioides* and the non-pathogenic related species *Uncinocarpus reesii* ^16,17^, as well as diverse dermatophytes that cause skin infections ^18,19^. Recently, the genomes of four non-pathogenic Onygenaceae species closely related to *Coccidioides* were described, providing additional resolution into changes in gene content between pathogenic and non-pathogenic Onygenales ^20^. Together these studies helped establish how gene content is related to the life cycle of different dimorphic pathogenic fungi and dermatophytes; for example, gene family contractions in cellulases and other plant metabolism genes, and gene family expansions in proteases, keratinases and other animal tissue metabolism genes, indicated that dimorphic fungi switched from a nutrition system based on plants to a system based on animals, though mostly relative to outgroups to the Onygenales such as *Aspergillus* ^14,17–19^. In the Onygenales, this hypothesis was tested experimentally; comparing growth on a wide variety of compounds revealed that in *Uncinocarpus reesii* hyphal growth was restricted on carbohydrates and considerably improved on proteins ^14^.

To examine key transition points in evolution of virulence and host adaptation in the dimorphic fungi, we increased the phylogenetic density within the Ajellomycetaceae by sequencing the genomes of non-pathogenic and emerging species and then performed comparative genomic analyses of the systemic, opportunistic, and non-pathogenic species. Although representative genomes for most of the described pathogenic Onygenales genera have been sequenced, the only strictly non-pathogenic species that have been sequenced are closely related to *Coccidioides* in the family Onygenaceae. Here, we have sequenced four additional Onygenales species from the family Ajellomycetaceae; this includes *H. griseus* and *P. hystricis*, neither of which has been associated with disease in mammals, and two additional adiaspore-forming strains of *Ea. parva* and *Ea. crescens* with distinct phenotypic properties. Leveraging our previous genomic studies and these additional genome sequences, we have characterized the extent of gene family expansions and contractions to provide a better understanding of the evolutionary mechanisms of adaptation in systemic dimorphic fungi in the Ajellomycetaceae.

## RESULTS

### Genome comparisons of Ajellomycetaceae, the largest group of dimorphic human pathogenic fungi

Building on previous genomic analyses of the members from the Ajellomycetaceae ^5,9,14,15,21^, we sequenced using Illumina technology the genomes of four species within Ajellomycetaceae with demarcated phenotypic differences. In addition to *Emmonsia parva* UAMH139 and *Emmonsia crescens* UAMH3008, we sequenced the genomes of one strain of *Emmonsia parva* UAMH130 (type strain) and one additional strain of *E. crescens* UAMH4076. *E. parva* UAMH130 is from lungs of a rodent in USA, and *E. crescens* UAMH4076 (type strain) is from a greenhouse source in Canada ^22^. We also sequenced two species that live in soil, where they are found in animal excrement but not associated with disease; neither species appears to undergo a thermal dimorphic switch to a pathogenic phase, as there is no observation of a yeast-like form ^13,22^. This includes one strain of *Helicocarpus griseus*, UAMH5409 (type strain; synonym *Spiromastix grisea*), isolated from gazelle dung in Algeria, and one strain of *Polytolypa hystricis*, UAMH7299, isolated from porcupine dung in Canada.

The *de novo* genome assemblies of these four species have similar sizes to those of other genera within the Ajellomycetaceae including *Histoplasma*, *Emergomyces* and *Paracoccidioides* ^5,9,14,15^, and have high representation of conserved eukaryotic genes (**Figure S1, S2**). The assembly sizes were 27.8 Mb in *E. parva* UAMH130, 33.8 Mb in *E. crescens* UAMH4076, 32.0 Mb in *P. hystricis*, and 34.7 Mb in *H. griseus* (**Table 1**). In addition, using both the CEGMA ^23^ and BUSCO ^24^ conserved eukaryotic gene sets, we found that the predicted genes for each sequenced species are nearly complete; all of these genomes include 96-98% of the core genes (**Figure S2**). The total numbers of predicted genes in *E. parva* UAMH130 (*n* = 8,202), and *E. crescens* UAMH4076 (*n* = 8,909) were similar to those found in annotated genome assemblies of other Ajellomycetaceae (*n* = 9,041 on average); however, relative to other Ajellomycetaceae *P. hystricis*, and *H. griseus* have a modest increase in gene content (*n* = 9,935 and *n* = 10,225, respectively; **Table 1**; **Figure S1**).

**Table 1.** Statistics of the annotated genome assemblies of *Emmonsia*, *Helicocarpus* and *Polytolypa*

| Species | Strain | Assembly (Mb) | Scaffolds (count) | N50 (kb) | GC (%) | Repeat (%) | Genes (count) |
| --- | --- | --- | --- | --- | --- | --- | --- |
| *Emmonsia parva* | UAMH130 | 27.8 | 383 | 173.2 | 47.5 | 2.6 | 8,202 |
| *Emmonsia crescens* | UAMH4076 | 33.8 | 1,288 | 96.6 | 42.9 | 15.0 | 8,909 |
| *Helicocarpus griseus* | UAMH5409 | 34.7 | 641 | 135.7 | 48.0 | 1.8 | 10,225 |
| *Polytolypa hystricis* | UAMH7299 | 32.0 | 546 | 124.6 | 45.6 | 12.8 | 9,935 |

Examining the guanine-cytosine (GC) content in the genomes *E. parva* UAMH130, *E. crescens* UAMH4076, *H. griseus* UAMH5409 and *P. hystricis* UAMH7299 revealed that none of these genome assemblies showed evidence of the pronounced bimodality of GC-content observed in *B. gilchristii* and *B. dermatitidis* (**Figure S3**; **Table 1**; ^5^). This highlights that the acquisition of repetitive elements contributing to bimodal GC distribution were a unique phenomenon during the evolution of the *B. gilchristii* and *B. dermatitidis* within the Onygenales. In addition, UAMH139 showed a small peak of low GC that is largely absent in UAHM130 (44.4% and 47.5% GC, respectively; **Table 1**), which is consistent with the phylogenetic divergence between these two strains, and highlights an independent gain of repetitive sequences (10.4% and 2.6% repetitive bases in UAMH139 and UAMH130, respectively; **Table 1**). Similarly, *E. crescens* UAMH3008 and UAHM4076 have demarcated differences in GC and repetitive content; UAHM4076 has a higher peak of low GC sequences (42.9% GC, 15.0% repetitive bases in UAMH4076; 45.2% GC, 5.1% repetitive bases in UAMH3008; **Table 1**). Additionally, the more basal strain *P. hystricis* UAMH7299 has a representation of low GC sequences (45.6% GC, 12.8% repeat), but in *H. griseus* UAMH5409 they are almost absent (48.0% GC, 1.8% repeats; **Figure S3**). These results highlight the independent expansion of repetitive elements during Ajellomycetaceae evolution.

### Phylogenomic analysis revealed multiple transitions within dimorphic pathogens to enable human infection

We estimated a strongly supported phylogeny of the newly sequenced *E. parva* UAMH130, *E. crescens* UAMH4075, *H. griseus* and *P. hystricis* strains relative to other dimorphic fungi using maximum likelihood and 2,505 single copy core genes (**Figure 1**). Branch length and topology supported the placement and relationships of *P. hystricis* and *H. griseus* within the Ajellomycetaceae as basally branching. The two *E. parva* isolates (UAMH130 and UAMH139) do not form a single well-defined clade, confirming that *E. parva* may not be a single species. In addition to longer branch lengths suggesting higher divergence of these isolates, the average genome-wide identity between UAMH130 and UAMH139 is much lower even comparing across groups that represent recently subdivided species, for example, within *B. gilchristii* and *B. dermatitidis* or *P. lutzii* and *P. brasiliensis*, with an average identity of 88.6% versus 95.8% and 91.1% respectively (**Figure S4**). *E. parva* (type strain UAMH130) is the most basal member of the clade including *E. parva* UAMH139 and *Blastomyces* species *B. percursus*, *B. gilchristii*, and *B. dermatitidis*. Previous molecular approaches ^12,25^ and recent multi-locus approaches ^8,9^ also support the polyphyletic nature of *E. parva* isolates. Although both isolates of *E. crescens* (UAMH3008 and UAMH4076) form a single clade, the genetic variation between them is also greater than that observed within other species from the same genus in the Ajellomycetaceae, with long branch lengths and slightly lower average genome identity (91.8%) between these isolates than seen in other intraspecies comparisons within the family (**Figure S4**). The *E. crescens* clade is closely related to the recently proposed genus *Emergomyces* including the *E. pasteurianus/africanus* species ^9^. The *E. crescens*-*Emergomyces* clade is a sister group of the clade including *Histoplasma* and *Blastomyces*, and *Paracoccidioides* is in a basal position for the dimorphic pathogens of the Ajellomycetaceae (**Figure 1**).

**Figure 1.**
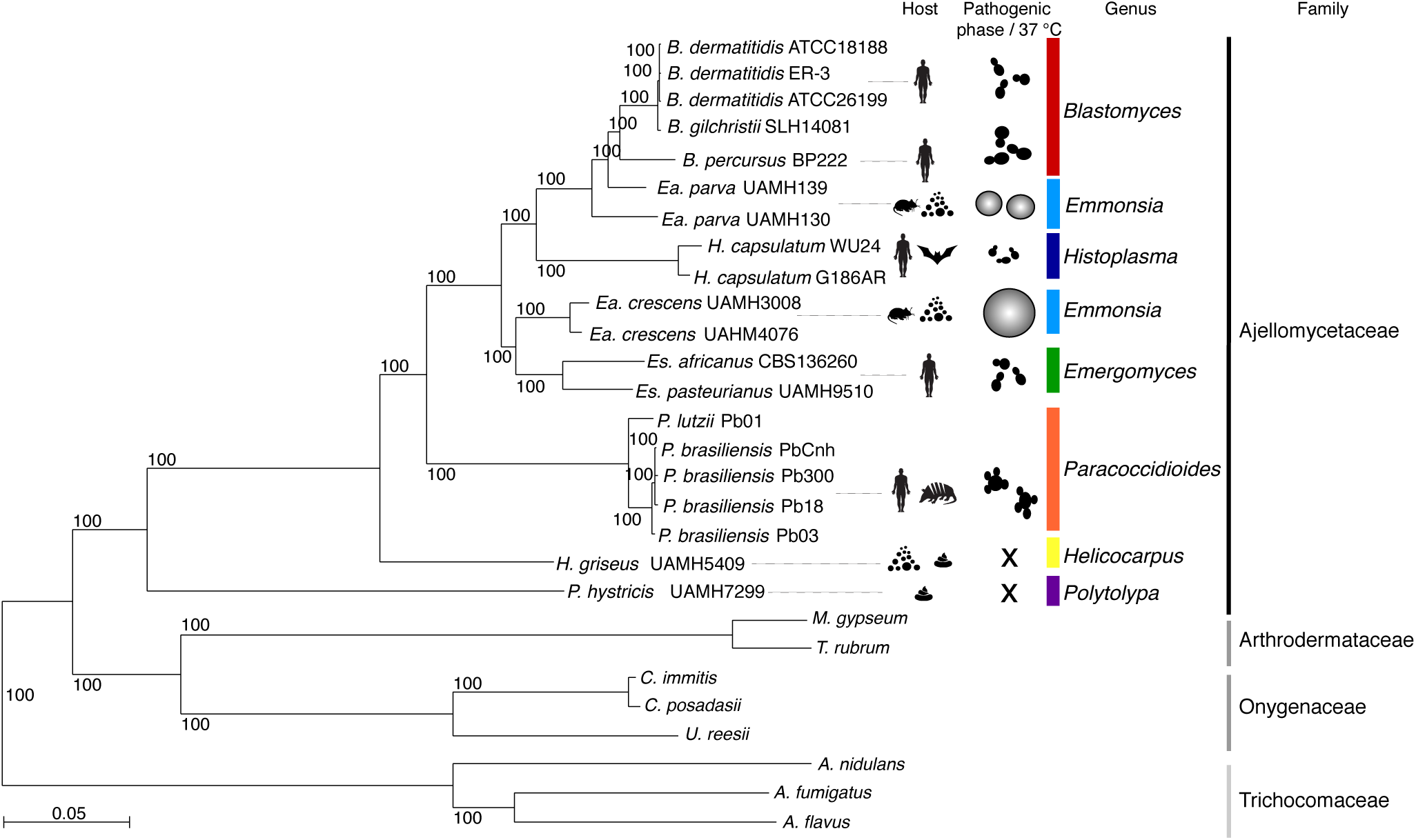
Phylogenomic relationships and diversity in the family Ajellomycetaceae. Maximum likelihood phylogeny (2,505 core genes based on 1,000 replicates) including 20 annotated genome assemblies of species within the family Ajellomycetaceae, five other Onygenales and three *Aspergilli*. All nodes were supported by 100% of bootstrap replicates. Color bars indicate the genus and icons represent most common source or host. Illustrations of the most common morphology at 37 °C or in the host are depicted for selected species highlighting the phenotypic variation within the Ajellomycetaceae and the transition to human hosts represented by pathogenicity traits.

The intercalated clades of primary pathogens (e.g. *H. capsulatum*, *P. brasiliensis* and *B. dermatitidis*), opportunistic pathogens (e.g. *E. pasteurianus*, *E. africanus*) producing a parasitic yeast phase, and non-pathogens producing adiaspore-type cells rather than yeast cells (e.g. *E. parva* and *E. crescens*) suggests that the dimorphic fungi in the Ajellomycetaceae have undergone numerous evolutionary transitions allowing adaptation, infection and virulence to humans and other mammals, and that interactions with other eukaryotes in the environment may help maintain the capacity for pathogenic growth in a mammalian host (**Figure 1**; **Table 2**). The non-thermally-dimorphic species *H. griseus* and *P. hystricis* are highly supported as early diverging lineages of Ajellomycetaceae as previously suggested ^13^; genomic analysis strongly supports *P. hystricis* as the earliest diverging lineage within the sequenced Ajellomycetaceae, and large genetic variation between these early saprophytes. The basal position of both *H. griseus* and *P. hystricis* in the family Ajellomycetaceae and the large divergence between them suggests that the history of this family could involve evolution from saprophytic non-thermotolerant and non-yeast-forming ancestral species to dimorphic human pathogenic species after the divergence with *H. griseus*, including the adaptation to higher temperatures and the appearance of a dimorphic switch and a yeast parasitic phase (**Figure 1**).

**Table 2.** Thermo-dimorphic pathogenic fungi, expansion and diversity within Ajellomycetaceae

| Species | Family | Dimorphism | Parasitic phase | Virulence in human |
| --- | --- | --- | --- | --- |
| *Blastomyces gilchristii* | Ajellomycetaceae | Yes | Intermediate yeast | Primary pathogen |
| *Blastomyces dermatitidis* | Ajellomycetaceae | Yes | Intermediate yeast | Primary pathogen |
| *Blastomyces percursus* | Ajellomycetaceae | Yes | Large yeast | Primary pathogen |
| *Emmonsia parva* | Ajellomycetaceae | Yes | Small adiaspore | Occasional |
| *Histoplasma capsulatum* | Ajellomycetaceae | Yes | Small yeast | Primary pathogen |
| *Emmonsia crescens* | Ajellomycetaceae | Yes | Large adiaspore | Occasional |
| *Emergomyces pasteurianus* | Ajellomycetaceae | Yes | Small yeast | Opportunistic |
| *Emergomyces africanus* | Ajellomycetaceae | Yes | Small yeast | Opportunistic |
| *Paracoccidioides brasiliensis* | Ajellomycetaceae | Yes | Multibudding yeast | Primary pathogen |
| *Paracoccidioides lutzii* | Ajellomycetaceae | Yes | Multibudding yeast | Primary pathogen |
| *Lacazia loboi* | Ajellomycetaceae | ND | Multibudding yeast | Primary pathogen |
| *Helicocarpus griseus* | Ajellomycetaceae | No | No | Non-pathogen |
| *Polytolypa hystricis* | Ajellomycetaceae | No | No | Non-pathogen |
| *Coccidioides immitis* | Onygenaceae | Yes | Spherules | Primary pathogen |
| *Coccidioides posadasii* | Onygenaceae | Yes | Spherules | Primary pathogen |
| *Sporothrix schenckii* | Ophiostomataceae | Yes | Small yeast | Primary pathogen |
| *Talaromyces marneffeii* | Trichocomaceae | Yes | Small septate yeast | Primary pathogen |

### Substantial gene family contractions underlie the shift from growth on plant to animals

By comparing the gene conservation and functional annotation of primary and opportunistic dimorphic pathogens with their saprophytic relatives *H. griseus* and *P. hystricis*, we identified significant decreases in gene family size (PFAM domain and GO-term counts, corrected p-value < 0.05) associated with a host/substrate shift from plants to animals in the pathogenic Ajellomycetaceae (**Figure 2**). Plant cell wall degrading enzymes are reduced in number or totally absent in dimorphic pathogens compared with *H. griseus* and *P. hystricis*, including several glycosyl hydrolases, and fungal cellulose binding domain-containing proteins (**Figure 2; Table S2**). These gene family changes suggest that dimorphic fungi from Ajellomycetaceae are not typical soil fungi in that they might maintain a close association with living mammals or other organisms such as protozoa or insects. This transition appears to have occurred within the family Ajellomycetaceae after divergence from *Helicocarpus* and *Polytolypa* (**Figure 1**). Our analysis also recapitulates previous reports comparing smaller numbers of genomes from the order Onygenales ^5,14,17,20^ that showed that genes coding for enzymes involved in the deconstruction of plant cell walls were absent from all the human pathogens (e.g. *Paracoccidioides*, *Blastomyces*, *Histoplasma*, *Coccidioides*) and non-pathogenic Onygenaceae (e.g. *Uncinocarpus reesii*, *Byssoonygena ceratinophila*, *Amauroascus mutatus*, *Amauroascus niger*, *Chrysosporium queenslandicum*) whereas these enzymes were commonly conserved in the non-pathogens outside the order Onygenales (including *Aspergillus, Fusarium*, *Neurospora*; **Figure 2; Table S2**). Notably, increasing the phylogenetic density within the order Onygenales in this study revealed that the loss of plant cell wall degradative enzymes occurred independently in two families within the Onygenales, and this highlights transitions between saprophytic and pathogenic species, where *Helicocarpus* and *Polytolypa* have retained enzymes required to live on plant material in the soil, whereas non-pathogenic species from the Onygenaceae, which are not adapted to survive in mammals, have also lost capacity to degrade plant material.

**Figure 2.**
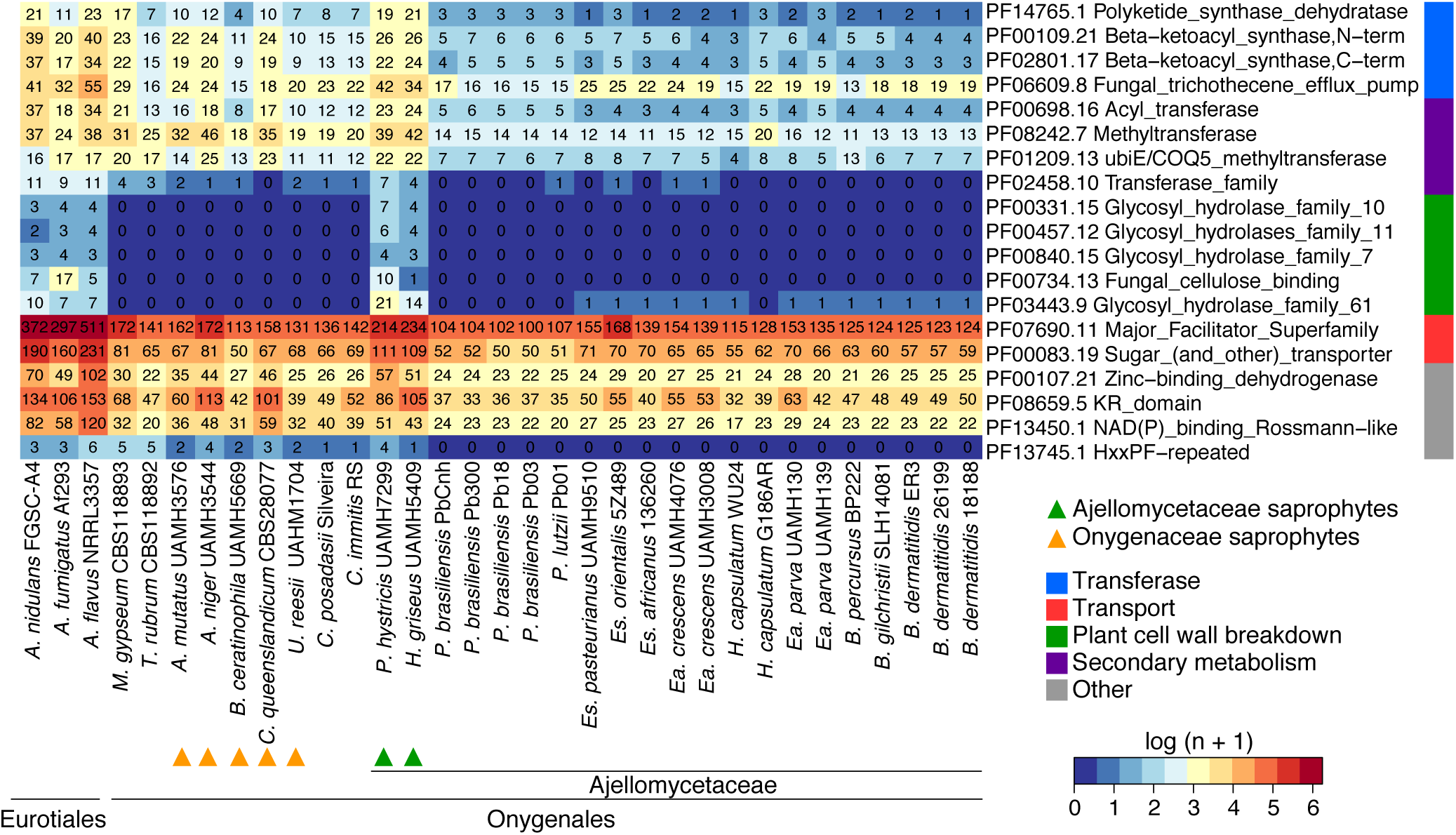
Gene family changes across the Ajellomycetaceae. Heatmap depicting results of a protein family enrichment analysis (PFAM domains; corrected *p-value <* 0.05) comparing the gene content of the non-dimorphic/non-pathogenic *Polytolypa hystricis* and *Helicocarpus griseus* with all other depicted members of the family Ajellomycetaceae including the dimorphic pathogens *Histoplasma*, *Emmonsia*, *Blastomyces*, *Emergomyces* and *Paracoccidioides*. Values are colored along a blue (low counts) to red (high counts) color scale, with color scaling relative to the low and high values of each row. Each protein family domain has a color code (right) indicating a global functional category. *Es*: *Emergomyces*; *Ea*: *Emmonsia*.

Other notable changes in gene content included loss of protein families associated with secondary metabolite biosynthesis such as polyketide synthase dehydratases, and beta-ketoacyl synthases domains in dimorphic Ajellomycetaceae relative to *H. griseus* and *P. hystricis* (**Figure 2**; **Table S3**). We identified predicted biosynthetic clusters using the program antiSMASH (antibiotics and Secondary Metabolism Analysis Shell ^26^). Overall, dimorphic pathogenic fungi from Ajellomycetaceae have fewer polyketide synthase (PKs) gene clusters than the saprophytic *H. griseus* and *P. hystricis* (**Table S3; Figure S5**). Dimorphic pathogens had lost essential genes or complete clusters of the type 1 and type 3 PKs, and terpene clusters (**Table S3**). Type 3 PKs are not common in fungi, and were not previously reported in the Ajellomycetaceae. Furthermore, *H. griseus* and *P. hystricis* but not dimorphic fungi have genes related to those involved in antibiotic biosynthetic pathways (ko01055; **Table S2**). Previous studies had shown that *P. hystricis* secretes a pentacyclic triterpenoid exhibiting antifungal and antibiotic activity, denominated Polytolypin ^27^, suggesting this terpene cluster is intact and may produce this molecule. Loss of such secondary metabolic pathways may weaken responses to other microbes in the environment. In addition to secondary metabolite biosynthesis, Ajellomycetaceae dimorphic fungi showed significant decreases in pathways related with the production of sphingolipids, chloroalkane and chloroalkene degradation, linoleic acid metabolism, and bisphenol degradation (KEGG-EC counts, corrected p-value < 0.05; **Table S2**). Loss of degradative pathways for bisphenol and chloralkanes in particular likely reflect encountering these molecules in the environment but not during pathogenic growth. Altogether, these shifts in gene content highlight how the Ajellomycetaceae dimorphic pathogenic fungi became less adapted to survive in the soil than the more basally diverging saprophytic species *H. griseus* and *P. hystricis*.

### Transition among carbohydrate metabolism and protein catabolism

We found that the evolution of carbohydrate metabolism and protein catabolism also occurred independently within the Ajellomycetaceae. We annotated carbohydrate active enzymes (CAZy) and peptidases (MEROPS; **Methods**), and using enrichment analysis found substantial shifts in the relative proportion of these groups among dimorphic and saprophytic species. Notably, the pathogenic Ajellomycetaceae species, but not the saprophytic species *H. griseus* and *P. hystricis*, showed a dramatic reduction of carbohydrate active enzymes, including 44 categories that were totally absent and 29 that were significantly depleted; these included glycoside hydrolases, glycosyltransferases, carbohydrate esterases, and polysaccharide lyases (corrected p < 0.05; **Figure 3**). The fact that these enzymes are reduced in dimorphic pathogenic Ajellomycetaceae but not in *H. griseus* and *P. hystricis* confirms that reduced carbohydrate metabolism occurred independently Ajellomycetaceae from the reduction in the Onygenaceae, originally noted in the spherule-dimorphic pathogen *Coccidioides* ^17^. In contrast to carbohydrate active enzymes, we found that the total number of peptidases is relatively similar in dimorphic and saprophytic species within the Ajellomycetaceae. Significant protein family expansions were found to be specific in *Blastomyces*, *Histoplasma* and Onygenaceae and Arthrodermataceae species relative to *P. hystricis* and *H. griseus*, including the A11, S12, and S08A peptidases (corrected p < 0.05; **Figure 3**). The peptidase family A11 contains endopeptidases involved in the processing of polyproteins encoded by retrotransposons, and the increase in copy number is correlated with the genome expansion due to the proliferation of repetitive elements in *Blastomyces* ^5^. The peptidase family S12 contains serine-type D-Ala-D-Ala carboxypeptidases, and S8 contains the serine endopeptidase subtilisin. Two families of fungal metalloendopeptidases (S8 and M35) were previously reported to be expanded in *Coccidioides* relative to saprophytic species outside the order Onygenales ^17^. In addition, it has been reported that the Ajellomycetaceae includes fewer copies of multiple classes of peptidases (M36, M43, S8) as well as an associated inhibitor (I9, inhibitor of S8 protease) relative to other Onygenales ^5^. Comparing across all the Onygenales, we found that all pathogenic Onygenales, as well as saprophytes related to *Coccidioides* such as *U. reesii*, had a higher ratio of proteases to carbohydrate active enzymes than the saprophytic Ajellomycetaceae *H. griseus* and *P. hystricis* (**Figure 3**). This strengthens the evidence that the adaptation and specialization of dimorphic Ajellomycetaceae occurred independently and that non-pathogenic Onygenaceae may represent an intermediate state in showing gene content changes associated with pathogens, which allowed these fungi to use proteins as a main source of energy in nutrient-limited environments.

**Figure 3.**
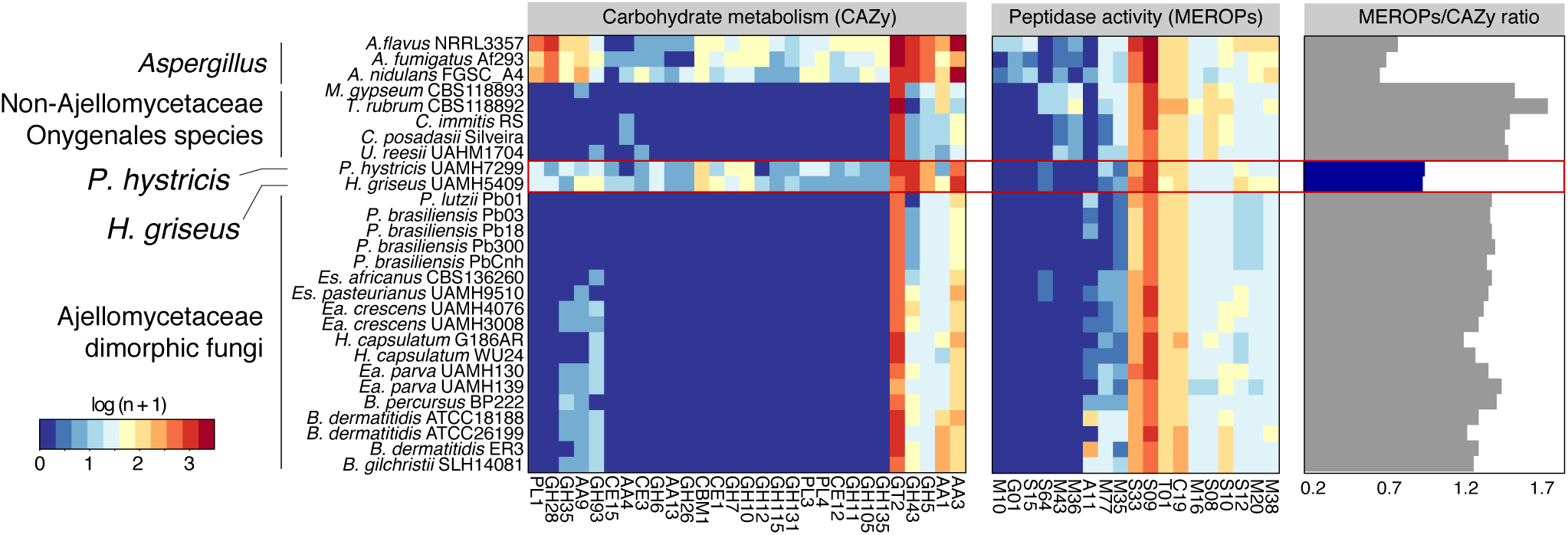
Shifts in carbohydrate metabolism and peptidase activity in Ajellomycetaceae. Heatmaps depicting for each taxon carbohydrate metabolism categories (CAZy) and peptidase families (MEROPs) across the Ajellomycetaceae species included in this study and other compared genomes. These categories were found significantly enriched (corrected *p-value <* 0.05) in the non-dimorphic systemic fungi *P. hystricis* and *H. griseus* relative to Ajellomycetaceae dimorphic fungi, or within Ajellomycetaceae species and other compared genomes from Onygenaceae and Arthrodermataceae families, or Eurotiales order. The ratio of MEROPS genes to CAZY genes for each genome across the Ajellomycetaceae species included in this study and other compared genomes is shown at right. *Es*: *Emergomyces*; *Ea*: *Emmonsia*.

### Loss of transporters in Ajellomycetaceae dimorphic fungi

In addition to major shifts in enzymes that support nutrient acquisition, the dimorphic Ajellomycetaceae have undergone losses of families of transporters. These include the large classes of major facilitator superfamily transporters (PF07690) and sugar transporters (PF00083) (**Figure 2)**. The number of transporters correlates with the numbers of genes predicted to have transmembrane helices (1,848 and 1,745 proteins have predicted transmembrane domains in *H. griseus* and *P. hystricis*, respectively, relative to an average of 1,490 in dimorphic Ajellomycetaceae pathogens, **Figure 4**). To characterize gene gain and losses in transporters, we annotated transporter families (**Methods**, ^28^), and found significant expansion of 18 classes of transporters encompassing 9 families in *H. griseus* and *P. hystricis* relative to dimorphic Ajellomycetaceae pathogens (corrected *p-value <* 0.05; **Figure 4**). Transporter families that were totally absent or significantly reduced in pathogenic species included the sugar porter, anion:cation symporter, AAA-ATPase, ammonium transporter and carnitine O-Acyl transferase. The substrates of these transporters include galacturonate, Rip1, hexoses, dipeptides, allantoate, ureidosuccinate, allantoin, glycerol, sulfate, sulfite, thiosulfate, sulfonates, tartrate, thiamine, cellobiose, cellodextrin, ammonia, methylamine, phenylacetate, beta-lactams, trichothecene, and carnitine O-octanoyl (**Figure 4**). Most of these compounds are only present in the plant cell wall, including the cellulose, hemicellulose, or pectin types, supporting that dimorphic pathogens have evolved to have more limited compounds from plant cell wall as source of energy.

**Figure 4.**
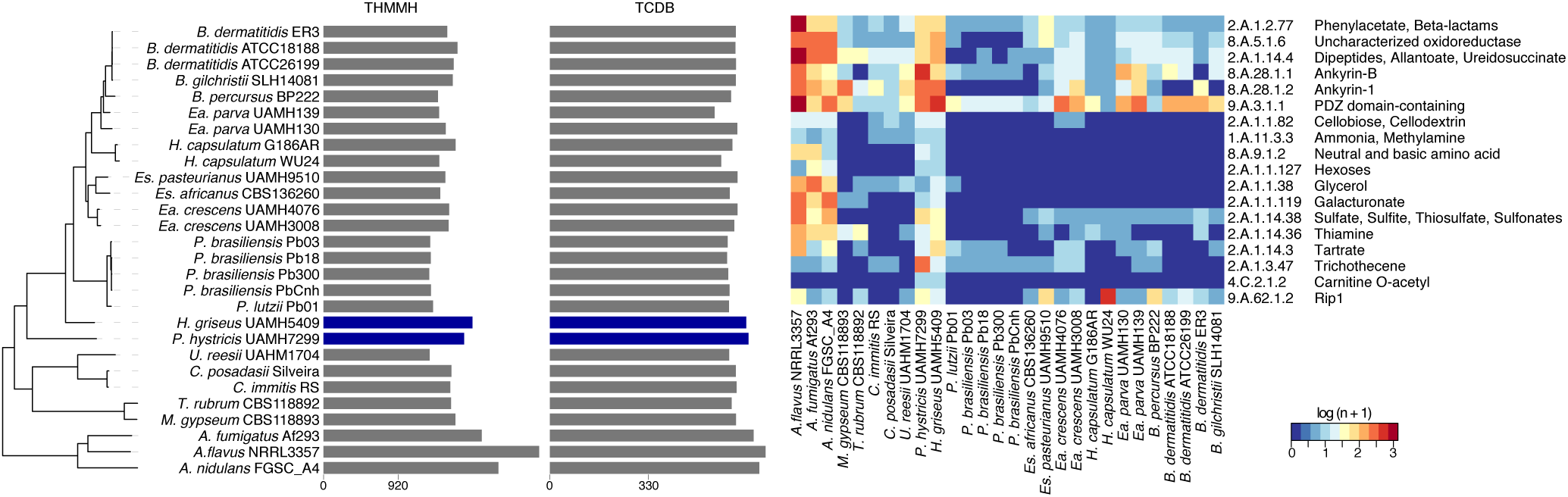
Shifts in metabolism matched substrate preference within Ajellomycetaceae contraction of transporters within the Ajellomycetaceae. (right) Heatmap depicting the number of family transporters (TCDB; color-code blue: low, red: high) for each taxon across the Ajellomycetaceae species and other compared genomes. These families were found significantly enriched (test, corrected *p-value <* 0.05) in the non-dimorphic *P. hystricis* and *H. griseus* relative to dimorphic Ajellomycetaceae fungi. (left) Total number of transporters identified in the genomes for each taxon included in this study. *Es*: *Emergomyces*; *Ea*: *Emmonsia*.

### Pathogenesis-related genes are conserved in both saprophytic and pathogenic lifestyles

Genes known to be important in the response to stress imposed by the host, including virulence-associated or yeast-phase specific genes of central importance in dimorphic fungi in the Onygenales, are conserved in *H. griseus* and *P. hystricis*. These include heat shock response proteins (*HSF*, *HSP90*, *HSP70*), dimorphic switch related proteins (*RYP1*, *RYP2*, *RYP3*, *AGS1*, *FKS1*), oxidative stress and hypoxia response proteins (*CATB*, *CATP*, *SOD3*, *SRB1*), as well as antigens (*PbGP43*, *PbP27*, *BAD1*; **Table S4;**^21^). In dimorphic pathogenic fungi both morphological transitions and growth temperature are linked, and loss of genes necessary for high-temperature growth and morphological transition in these pathogens results in attenuated virulence, which highlights that they are essential for pathogenesis ^29–31^. Some of these genes such as the *RYP*1-3 transcriptional regulators and the dimorphism-regulating histidine kinase *DRK1* have been connected with morphogenetic adaptation in response to environmental stimuli, and with key determinants of the mycelia to yeast transition in dimorphic human pathogenic fungi in the Ajellomycetaceae ^29,30^. Although *H. griseus* and *P. hystricis* are not dimorphic and cannot grow at 37 °C, the heat shock response has been shown to be highly evolutionarily conserved among eukaryotes. The fact that genes of central importance for high-temperature growth and dimorphic switch are conserved in *H. griseus* and *P. hystricis* suggests that these saprophytes may sense different gradations of temperature, or that the mechanism that triggers a rapid response to the adaptation may not be rapid enough to enable these fungi to grow at 37 °C.

Dimorphic Ajellomycetaceae may have evolved specialized mechanisms to regulate their transcriptional responses; these include expansion of proteins related to the regulation of transcription, and protein kinase activity (GO term enrichment analysis; *q*-value < 0.05; **Table S2**), which are all expanded in dimorphic pathogens relative to *H. griseus* and *P. hystricis*. Thus, the gain of transcription factors, transcriptional regulators and phosphotransferases suggests that rapid evolution of transcriptional mechanisms may underlie the adaptation to both high temperature and the dimorphic transition. As one of the largest categories enriched in dimorphic Ajellomycetaceae but not in saprophytic species was the protein kinase activity (**Table S2**), we further classified protein kinases using Kinannote ^32^, including the divergent fungal-specific protein kinase (*FunK1*; **Methods**). We found an expansion of the *FunK1* family, in dimorphic Ajellomycetaceae as previously reported ^5,14^. As we increased the phylogenetic density in the Ajellomycetaceae, including species and strains with high genetic and phenotypic variation this allowed resolution of the evolutionary history of this family of kinases (**Figure 5**). *FunK1* family is enriched in the systemic pathogens *B. dermatitidis/gilchristii*, *H. capsulatum*, *Paracoccidioides*, but limited in saprophytic species, *H. griseus* and *P.* hystricis. Notably, this family is also limited in the adiaspore-forming *E. parva* (UAMH130, UAMH139), *E. crescens* (UAMH4076) but not in *E. crescens* (UAMH3008). In addition, we found that the recently described primary pathogen *B. percursus* includes a large expansion of FunK1 kinases, with almost twice the number as compared with its closest relative *B. dermatitidis* (**Figure 5**). Phylogenetic analysis of *FunK1* genes revealed that this family undergone independent lineage-specific expansions in primary dimorphic pathogens of both Ajellomycetaceae and Onygenaceae, but not in non-pathogenic or opportunistic dimorphic species (**Figure 5**).

**Figure 5.**
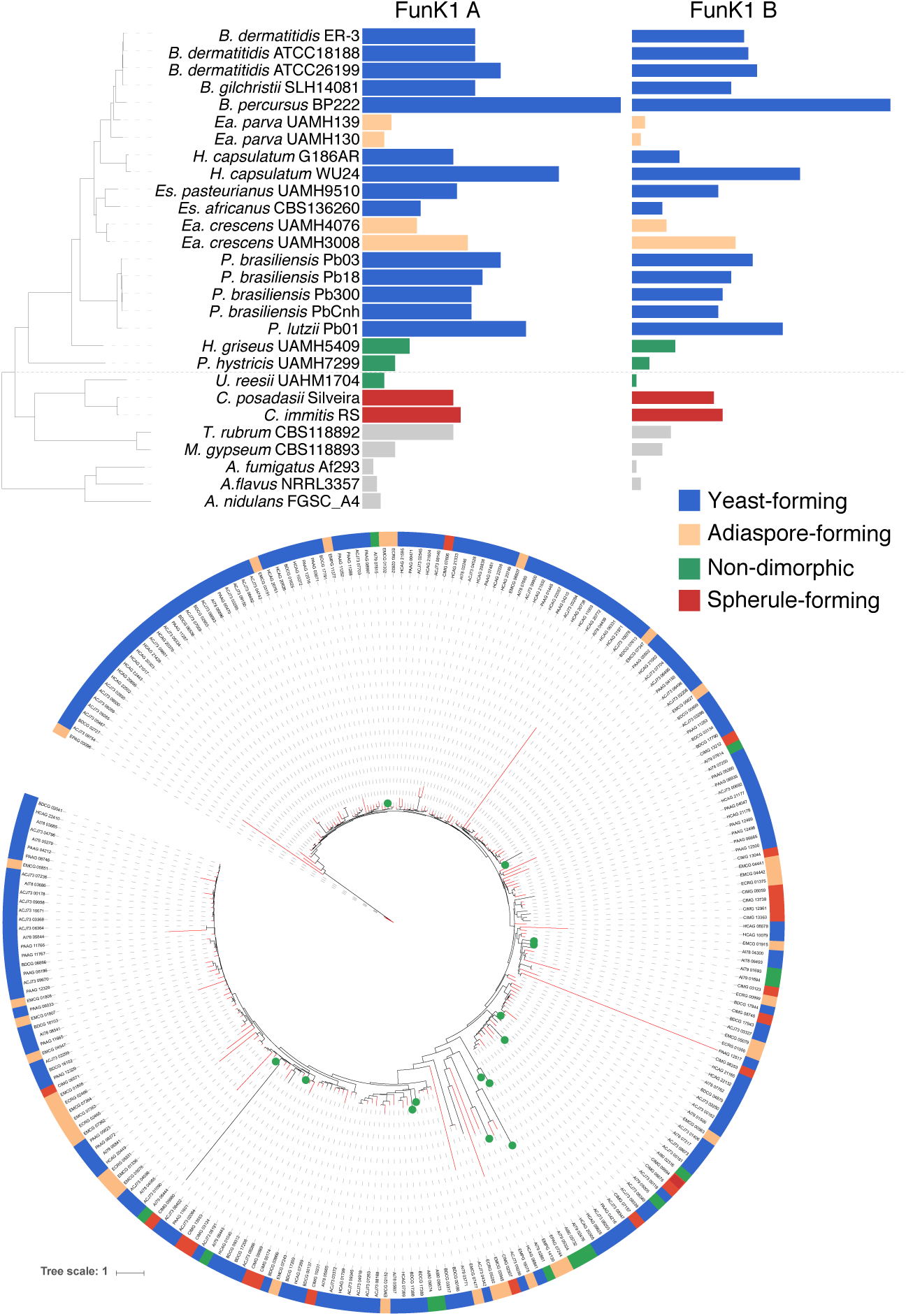
Expansion and contraction of protein kinases class *FunK1* across the Ajellomycetaceae. (top) *FunK1 A* and *FunK1 B* counts for each taxon across the Ajellomycetaceae species and other compared genomes. The tree shown corresponds to the maximum likelihood phylogenetic tree of the family Ajellomycetaceae from Fig. 1. (bottom) Maximum likelihood tree of *FunK1* family showing expansion in Ajellomycetaceae yeast-forming (blue) pathogenic species. Each species (top) or gene (bottom) has a color code indicating the parasitic phase at 37 °C (yeast, adiaspore, spherule, or non-pathogenic).

To identify specific genes that evolved within the Ajellomycetaceae to enable the dimorphic transition and infection of mammals, we identified and examined ortholog clusters that were unique to these pathogens and absent in both *H. griseus* and *P. hystricis*. We identified 75 ortholog clusters that were present in dimorphic Ajellomycetaceae and dimorphic Onygenaceae species but absent in *H. griseus* and *P. hystricis* (i.e. ‘sporophyte gene loss events’), and 212 ortholog clusters that were present in dimorphic Ajellomycetaceae, but absent in the dimorphic Onygenaceae species, *H. griseus* and *P. hystricis* (i.e. ‘dimorphic Ajellomycetaceae gene gain events’; **Table S5**). The gene loss events include a mold specific protein (CIMG_00805), a phenol 2-monooxygenase (BDFG_05966), as well as several protein kinases and transporters. The gene gain events include several protein kinases, transporters, transcription factors, and peptidases (**Table S5**). In addition, we annotated genes that have been broadly associated with host-pathogen interactions (Pathogen Host interaction database PHI), and genes that have been reported as virulence factors in fungal species (Database of Fungal Virulence Factors DFVF) ^33,34^. We found that many genes that have been associated with host-pathogen interactions are present in the saprophytic *P. hystricis* and *H. griseus*, and in fact some appear at higher copy number counts (**Table S5**). This emphasizes that host-pathogen interaction and virulence factors are not only important for pathogenicity in mammals, but also for saprophytic species to survive and adapt to stress in the environment, including for example the ability to evade the engulfment by amoebae, or the interaction with insects ^35,36^.

### Conservation of genes induced during *in vivo* infection

We also hypothesized that genes induced during mammalian infection in dimorphic Ajellomycetaceae pathogens might have different gene conservation patterns among *Blastomyces*, *Histoplasma*, *Paracoccidioides*, and the novel species and saprophytes. Transcriptional responses during an *in vivo* mouse model of infection had been studied in *B. dermatitidis* (strain ATCC26189; ^5^) and *P. brasiliensis* (strain Pb18; ^37^). In both *B. dermatitidis* and *P. brasiliensis* the high-affinity zinc transporter, *ZRT1* (BDFG_09159; PADG_06417) and the flavodoxin-like protein *PST2* (BDFG_08006; PADG_07749) were highly induced during infection; these genes are conserved in all Ajellomycetaceae included in this analysis (**Table S6**). Of the 72 *in vivo-*induced genes in *B. dermatitidis* ATCC26188, 22% are absent from the *P. hystricis* genome and 11% are absent from the *H. griseus* genome. The genes absent in *P. hystricis* include the cysteine dioxygenase (BDFG_08059), an acetyltransferase (BDFG_07903), a thioesterase (BDFG_02348), a superoxide dismutase (BDFG_07895), a sodium/hydrogen exchanger (BDFG_05427), and a response regulator protein (BDFG_01066). In addition, we identified three secreted proteins that were found induced during the interaction of *B. dermatitidis* with macrophages (BDFG_08689, BDFG_06057, BDFG_03876) ^5^ that are found conserved only in the dimorphic Ajellomycetaceae pathogens but not in *H. griseus*, *P. hystricis*, or outgroup species. These are strong candidates for effector proteins involved in host infection. Similar numbers of genes found induced during *in vivo* infection in *P. brasiliensis* were absent in *P. hystricis* and *H. griseus*, (23% and 17.5%, respectively; **Table S6**). Genes absent in both species include Gpr1 family protein (PADG_08695), isovaleryl-CoA dehydrogenase (PADG_05046), glutathione-dependent formaldehyde-activating (PADG_05832), predicted transmembrane protein (PADG_05761), and ergot alkaloid biosynthetic protein A (PADG_07739). Many of the genes that were absent in *P. hystricis* and *H. griseus*, were also absent in at least one of the dimorphic pathogens in the Ajellomycetaceae, highlighting species-specific gene gain events that may confer virulence specialization.

### Evolution of the mating type locus in the Onygenales family Ajellomycetaceae

To elucidate the evolution of the mating system in the Ajellomycetaceae, we identified and characterized the mating locus of *P. hystricis*, and *H. griseus*. Notably, these two species have both mating type idiomorphs *HMG* box (*MAT 1-2*) and alpha box (*MAT1-1*); these are therefore the only homothallic species identified so far within the Ajellomycetaceae (**Figure 6**). Comparative analysis showed that the locus is not expanded relative to other Ajellomycetaceae species (21 kb between the flanking genes *SLA2* to *APN2/COX13*), unlike the expansion that observed in the larger genomes of *B. dermatitidis* or *B. gilchristii* (~60 kb; ^38^). One difference is that a sequence inversion between *APN2* and *COX13* is uniquely observed in *P. hystricis* relative to all other sequenced Ajellomycetaceae species, including *H. griseus* and previously reported species ^5,15,38,39^. This inversion observed in *P. hystricis* (mating type idiomorph flanked by *SLA2* and *APN2/COX13*) is similar to the gene order in the Onygenaceae species *Coccidioides* spp. but not in the Arthrodermataceae species *M. gypseum* (*Nannizzia gypsea* ^40^) or *T. rubrum* (**Figure 6; Table S7**) suggesting that independent inversion events within the locus. In addition, we identified the mating type locus in non-pathogenic Onygenaceae, and found that *Amauroascus mutatus* and *Byssoonygena ceratinophila* have both mating type idiomorphs (**Table S7**). In *A. mutatus*, *HMG* box (*MAT 1-2*) and alpha box (*MAT1-1*) are linked in the same scaffold, however they are separated by 184 kb, including 65 protein-coding genes (**Figure 6**). The identification of homothallic mating loci in non-pathogenic species within both the Ajellomycetaceae and the Onygenaceae suggests that there have been multiple transitions between homothallic and heterothallic sexual states in dimorphic pathogens and non-pathogenic species in the Onygenales.

**Figure 6.**
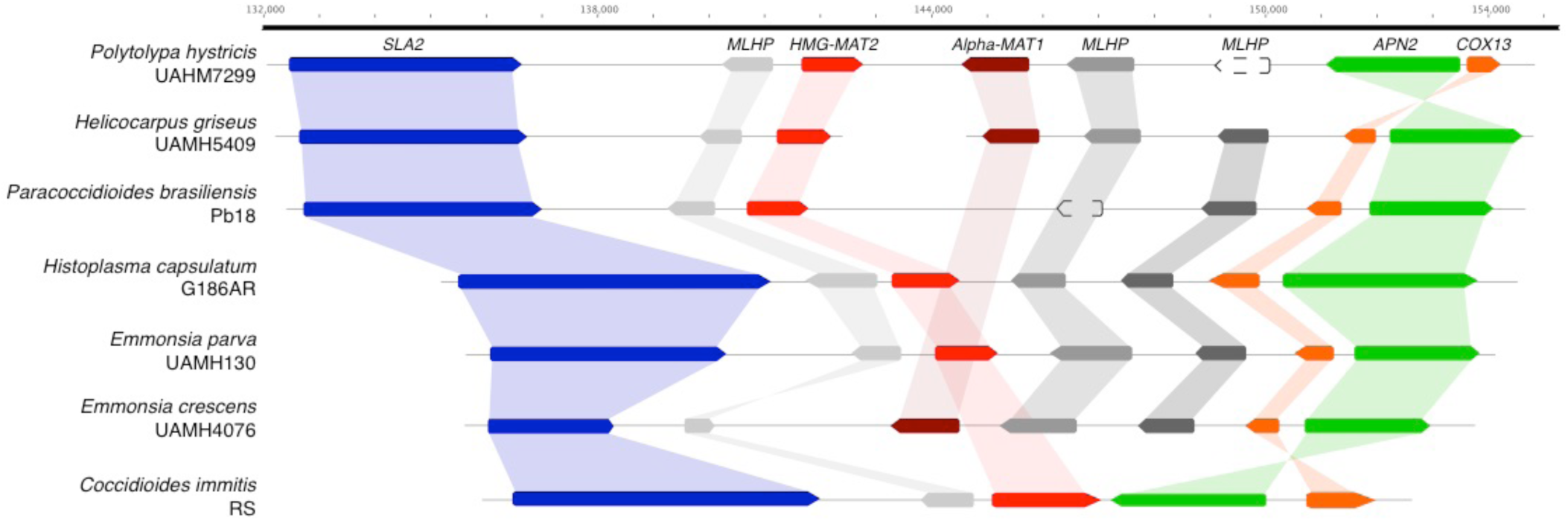
Mating type evolution within the Onygenales family Ajellomycetaceae. (top) Synteny schema depicting orientation and conservation of the genes adjacent to the mating type locus idiomorphs HMG box (*MAT 1-2*, red) and alpha box (*MAT1-1*, dark red) in the Ajellomycetaceae. The more basal species, *P. hystricis*, has both mating type idiomorphs intact, being the only homothallic species identified so far within the Ajellomycetaceae. The locus has similar size to other Ajellomycetaceae species with 21 kb average from *SLA2* (blue) to *APN2*/*COX13* (green/orange), but not *Blastomyces dermatitidis/gilchristii* (60 kb; Li W et al., 2013). A synteny inversion encompassing *APN2* and *COX13* is uniquely observed in this ancestor relative to the rest of Ajellomycetaceae species included in this study, including *H. griseus*. Grey genes are mating locus hypothetical proteins (MLHP).

## DISCUSSION

The evolution of the dimorphic Onygenales is characterized by multiple transitions between saprophytic and human pathogenic growth. By sequencing the genomes of two additional dimorphic adiaspore-forming species (*Emmonsia parva*, *E. crescens*) and two early diverging non-pathogenic, non-dimorphic species (*Helicocarpus griseus* and *Polytolypa hystricis)*, we analyzed how changes in gene content correlate with transitions to pathogenesis. Our results establish an outer bound on the timing of two key events in the evolution of the Ajellomycetaceae; both the loss of genes involved in digesting plant cell walls and the expansion of proteases, transcriptional regulators, and protein kinases, occurred after divergence with *H. griseus*. The shift in these families was previously observed in both *Paracoccidioides* within this group and in the related *Coccidioides*-*Uncinocarpus* clade ^14,17^; more recently the comparison with non-pathogenic Onygenaceae species supported previous reports of the contraction of gene families involved in digesting plant cell walls throughout the Onygenales, and of the expansion of gene families involved in digesting animal protein ^20^. Whereas the studies on contraction of gene families involved in digesting plant cell walls addressed other fungi outside the order Onygenales, our analysis shows that *H. griseus* and *P. hystricis* did not experience such a contraction, which suggests that contractions in gene families involved in digesting plant cell walls throughout the Onygenales have occurred multiple times independently and more recently in Onygenaceae and Ajellomycetaceae families.

Since many virulence factors and dimorphic switch related proteins were also conserved in the rarely pathogenic (*Ea. parva* and *Ea. crescens*) and in the non-dimorphic Ajellomycetaceae species (*P. hystricis* and *H. griseus*), this suggests that these proteins could be needed not only for survival in the host, and that presence of virulence factors alone does not specify pathogenicity. In dimorphic Ajellomycetaceae other mechanisms such as gene regulation or selection may play a role in how these genes respond during distinct stimuli. In addition, it is possible that those virulence factors in mammals were also needed for the survival in the soil, for example to survive interaction with amoebae or in insects ^41^. However, there are some virulence factors that are needed for virulence in mammals that are not necessary for virulence in other eukaryotes. For example, the alpha mating factor locus in *Cryptococcus neoformans* contributes to virulence in mice, but no difference was observed in the interaction of congenic *MATα* and *MAT***a** strains with amoebae ^42^. While the human host is a very different habitat in comparison with soil or excrements of animals that are the most common natural niche for this group of fungi, interactions with other eukaryotes in the environment likely selected for some properties that also predisposed species to become pathogenic to humans.

Selection pressures in the environment are responsible for the emergence and maintenance of traits that confer upon some soil microbes the capacity for survival in animal hosts. This is important to understand the evolution and emergence of novel pathogens. For example, *Emergomyces africanus* has recently emerged and numerous cases of novel yeast-forming species have been reported. However, *Emergomyces* species have not been found in other mammals like rodents or in the soil. In addition, *E. africanus* has a very restricted geographic distribution. Across the Onygenales, nearly all species have a high lower ratio of proteases to carbohydrate active enzymes; the exceptions are the two saprophytes *P. hystricis* and *H. griseus*, but not the saprophytes related to *Coccidioides* such as *Uncinocarpus reesii*. This genomic profile matches the results of growth assays for the non-pathogenic species *U. reesii*; this species can grow on a wide range of proteinaceous substrates but only a very limited range of carbohydrates, namely cellulose and its component glucose^14^. Together, these findings suggest that fungi in the Onygenales transitioned from saprophytes that digest plant materials to saprophytes that digest more limited plant materials and a wide range of animal proteins, and finally those that are major animal pathogens.

We found many of the genes with roles in the dimorphic transition and virulence are conserved in *Ea. parva* and *Ea. crescens*, and in saprophytic species *H. griseus* and *P. hystricis*, and we did not find larger patterns of gain of functional classes in pathogenic species. However, we found smaller changes in gene content related to the regulation of gene expression and signaling that may account for the ability to grow at high temperature, species specific morphological transitions, and therefore their differences in virulence, since rapid thermo-tolerance and morphological adaptation is essential for pathogenesis ^29,30,43^. Overall, we found significant expansions of the number of the protein kinase *FunK1* family, transcription factors and other genes associated with the regulation of gene expression in yeast-forming Ajellomycetaceae relative to adiaspore-forming and saprophytic species. Our comparison with saprophytic Ajellomycetaceae highlights dramatic decreases and complete absences of several classes of gene families that are important for survival and growth in a soil environment.

Intriguingly, we found that both more basal non-pathogenic Ajellomycetaceae species *H. griseus* and *P. hystricis* appear homothallic, in striking contrast to all the previously described largely pathogenic species in this group that are all heterothallic. *P. hystricis* has a fused mating type locus, with both idiomorphs, *HMG* box (*MAT 1-2*>) and alpha box (*MAT1-1*), fused at a single locus, and the same is likely true of *H. griseus*, as genes are located at the ends of two scaffolds in this assembly. In addition, we found that this is also true for the non-pathogenic Onygenaceae species *Amauroascus mutatus* and *Byssoonygena ceratinophila*; these species have both mating type idiomorphs. *A. mutatus* has both mating type idiomorphs located on the same scaffold; however, these are separated by a much larger distance than is typically observed at the *MAT* locus of homothallic fungi. This suggests that this locus has been subject to recombination events either to bring the two idiomorphs nearby on the same scaffold or alternatively in expansion of a fused locus. While it has been previously suggested that the ancestor of the Onygenales very likely was heterothallic ^39^, the placement of multiple lineages of homothallic species in this group raises the possibility of a heterothallic ancestor, with more recent transitions in pathogenic species to be homothallic and rearrangements of the mating locus in others such as *P. hystricis* and *A. mutatus*. Both *P. hystricis* UAMH7299 and *H. griseus* UAMH5409 can develop ascomata and ascospores, which have been shown to sporulate in *P. hystricis* ^11,27,44^. Loss of the capacity for homothallic self-mating could increase the frequency of out-crossing and this may be beneficial to some species under pressure from their environment or host ^45^ such as the pathogenic species in this group. Examining the mating type loci of additional non-pathogenic species in the Onygenales would further reveal if there are novel configurations of homothallic or heterothallic loci, which may help infer the mating ability of ancestral species and the timing of transitions in pathogens and non-pathogens.

Our phylogenomic analysis revealed that dimorphic pathogens in the Onygenales have undergone multiple evolutionary transitions that enabled infection of humans and other mammals. Some such as the loss of plant cell wall degrading enzymes are convergent for pathogens in the two families of Onygenales. Our comparisons also revealed major differences in non-pathogenic species between these two families, which suggested that some species appear intermediate between saprophytes and pathogens in terms of having a shift in metabolic capacity without associated pathogenic growth. Targeting additional intermediate species within the Ajellomycetaceae with different phenotypes (e.g. *Emmonsiellopsis*) for genome sequencing and comparison would help further refine the picture of the overall evolution of this important group of fungi.

## MATERIALS AND METHODS

### Genome sequencing

Species were selected for sequencing based on previous estimates of high genetic variation among the members of the genus *Emmonsia*, especially in the *E. parva* species, and based on observations of the emergence of *Emmonsia*-like species in the Ajellomycetaceae causing systemic human mycoses worldwide ^8,9^. We selected one strain of *Emmonsia parva* (UAMH130; CBS139881; type strain) and one additional strain of *Emmonsia crescens* (UAMH4076; CBS139868). *E. parva* UAMH130 is from the lungs of a rodent in the USA, and *E. crescens* UAMH4076 is from a greenhouse source in Canada, and mated with *E. crescens* UAMH129, 349 ^22^. In addition, the first strictly non-pathogenic non-dimorphic species from the Ajellomycetaceae, one strain of *Helicocarpus griseus* (UAMH5409; a.k.a. *Spiromastix grisea*) and *Polytolypa hystricis* (UAMH7299) were selected for whole genome sequencing. Genomic DNA was isolated from mycelial phase at 22 °C. We constructed one library with 180-base inserts and sequenced on the Illumina HiSeq 2000 platform to generate 101 bp paired-end reads with high coverage (173X UAMH4076; 199X UAMH130; 205X UAMH5409; 163X UAMH7299).

### Genome assembly, gene prediction and annotation

The 101-bp Illumina reads of *E. parva* UAMH130, *E. crescens* UAMH4076, *P. hystricis* and *H. griseus* were assembled using ALLPATHS-LG ^46^ with default parameters. All four *de novo* assemblies were evaluated using the GAEMR package (http://software.broadinstitute.org/software/gaemr/), which revealed no aberrant regions of coverage, GC content or contigs with sequence similarity suggestive of contamination. Scaffolds representing the mitochondrial genome were separated out from the nuclear assembly.

Genes were predicted and annotated by combining calls from multiple methods to obtain the best consensus model for a given locus. These included *ab initio* predictions (GlimmerHMM, Augustus, Snap, GeneMark-ES), homologous inference (Genewise, TBlastN), and gene model consolidation programs (EvidenceModeler) ^47,48^. For the protein coding-gene name assignment we combined HMMER PFAM/TIGRFAM, Swissprot and Kegg products. Kinannote was used to annotate protein kinases ^32^. To evaluate the completeness of predicted gene sets, the representation of core eukaryotic genes was analyzed using CEGMA genes ^23^ and BUSCO ^24^.

### Identification of orthologs and phylogenomic analysis

To compare gene content and conservation, we identified orthologous gene clusters in these sequenced genomes and in other Onygenales genomes using OrthoMCL (version 1.4) with a Markov in a on index of 1.5 and a maximum e-value of 1e-5 ^49^. We included the genomes of the classical dimorphic pathogenic species (*Blastomyces dermatitidis*, *B. gilchristii*, *Histoplasma capsulatum*, *Paracoccidioides brasiliensis*, *P. lutzii*, *Coccidioides immitis*, and *C. posadasii*), novel dimorphic human pathogenic species (*Blastomyces percursus*, *Emergomyces africanus*, *E. pasteurianus*), and two dimorphic non-human pathogenic species *Emmonsia parva* UAMH139 and *Emmonsia crescens*, along with other species, including three *Aspergillus* (**Table S1; Figure S7**). In addition, we annotated and included in the compare gene content and conservation analysis the available genomes of saprophytic species from the family Onygenaceae *Byssoonygena ceratinophila* (UAMH5669), *Amauroascus mutatus* (UAMH3576), *Amauroascus niger* (UAMH3544), and *Chrysosporium queenslandicum* (CBS280.77), and the genome of the yeast-forming dimorphic pathogen *Emergomyces orientalis* (5Z489, **Table S1**). Although, this is the largest genomic data set of Ajellomycetaceae strains so far, the number of shared gene families after each addition of genome data did not clearly reach a plateau; this suggests that the current estimate of a core of 4,200 genes is too high, and that more genomes are needed to define the core-genome and pan-genome in the Ajellomycetaceae (**Figure S6**).

To examine the phylogenetic relationship of the newly sequenced *E. parva* and *E. crescens*, *P. hystricis* and *H. griseus* relative to other dimorphic fungi we used single copy orthologs determined and clustered using OrthoMCL. A total of 31 genomes from the Onygenales order and three *Aspergillus* genomes were chosen to estimate the species phylogeny (**Table S1**). These include the four newly strains described here: *E. parva* UAMH130, *E. crescens* UAMH4076, *P. hystricis* UAMH7299 and *H. griseus* UAMH5409, as well as the following: three *Blastomyces dermatitidis* (ATCC26199, ATCC18188, ER-3), one *Blastomyces gilchristii* (SLH14081), one *Emmonsia parva* (UAMH139), one *Emmonsia crescens* (UAMH3008), two *Histoplasma* (WU24, G186AR), five *Paracoccidioides* (Pb01, Pb03, Pb18, PbCnh, Pb300), two *Coccidioides* (RS, Silveira), *Uncinocarpus reesii* (UAMH1704), *Microsporum gypseum* (CBS118893), *Trichophyton rubrum* (CBS118892), *Aspergillus nidulans* (FGSC A4), *A. flavus* (NRRL3357) and *A. fumigatus* (Af293). Protein sequences were aligned using MUSCLE, and a phylogeny was estimated from the concatenated alignments using RAxML v7.7.8 ^50^ with model PROTCATWAG with a total of 1,000 bootstrap replicates.

### Gene family and protein domain analysis

To identify gene content changes that could play a role in the evolution of the dimorphism and pathogenesis within Ajellomycetaceae, we searched for expansions or contractions in functionally classified genes compared to the other fungi from the order Onygenales. Genes were functionally annotated by assigning PFAM domains, GO terms, and KEGG classification. HMMER3 ^51^ was used to identify PFAM domains using release 27. GO terms were assigned using Blast2GO ^52^, with a minimum e-value of 1x10^−10^. Protein kinases were identified using Kinannote ^32^ and divergent FunK1 kinases were further identified using HMMER3. We further annotated carbohydrate active enzymes, peptidases, and transporter families using the CAZY version 07-15-2016 ^53^, MEROPS version 9.12 ^54^, and TCDB version 01-05-2017 ^28^ databases, respectively. Proteins were searched against their corresponding databases using BLAST, with minimum e-values of 1×10^−80^ for CAZY, 1×10^−20^ for MEROPS, and 1×10^−20^ for TCDB.

To identify functional enrichments in the compared genomes, we used four gene classifications: OrthoMCL similarity clusters, PFAM domains, KEGG pathways, and Gene Ontology (GO), including different hierarchy levels, MEROPS and CAZY categories, and transporter families. Using a matrix of gene class counts for each classification type, we identified enrichment comparing two subsets of queried genomes using Fisher’s exact test. Fisher’s exact test was used to detect enrichment of PFAM, KEGG, GO terms, CAZY, MEROPS, and transporter families between groups of interest, and p-values were corrected for multiple comparisons ^55^. Significant (corrected p-value < 0.05) gene class expansions or depletions were examined for different comparisons.

## Data Availability Statement

The raw sequences, genome assemblies and gene annotations were deposited in GenBank under the following BioProject accession numbers: *Emmonsia parva* strain UAMH130 (PRJNA252752), *Emmonsia crescens* strain UAMH4076 (PRJNA252751), *Polytolypa hystricis* strain UAMH7299 (PRJNA234736), and *Helicocarpus griseus* strain UAMH5409 (PRJNA234735).

## ACKNOWLEDGEMENTS

We thank Lynne Sigler for her help in selecting strains for sequencing. We thank the Broad Genomics Platform for generating Illumina sequence for this study, Sarah Young and Margaret Priest for their assistance in assembly and annotation, Christopher Desjardins for helpful comments on the manuscript, and Leslie Gaffney for help with Figure 1. This project has been funded in whole or in part with Federal funds from the National Institute of Allergy and Infectious Diseases, National Institutes of Health, Department of Health and Human Services, under award N°: U19AI110818. This work was partly supported by Colciencias via the grants under Contract N°: 122256934875 and N°: 221365842971, and by the Universidad de Antioquia via a “Sostenibilidad 2016” grant.

## Author contributions

Conceived and designed the experiments: JFM JGM OKC CAC. Performed the assembly and annotation: JFM CAC. Analyzed the data: JFM CAC. Contributed reagents/materials/analysis tools: JGM OKC. Wrote the paper: JFM CAC.

## Additional information

Competing interests: The authors declare no competing financial interests.

